# The Complex Ecosystem in Non Small Cell Lung Cancer Invasion

**DOI:** 10.1101/176446

**Authors:** Seth Haney, Jessica Konen, Adam I. Marcus, Maxim Bazhenov

**Affiliations:** Department of Medicine, Division of Pulmonary, Critical Care & Sleep Medicine, University of California, San Diego, La Jolla, CA; Department of Thoracic/Head and Neck Medical Oncology, The University of Texas MD Anderson Cancer Center, Houston, Texas, USA.; Winship Cancer Institute, Emory University, Atlanta, GA; Department of Hematology and Medical Oncology, Emory University, Atlanta, GA

## Abstract

Many tumors are characterized by genetic instability, producing an assortment of genetic variants of tumor cells called subclones. These tumors and their surrounding environments form complex multi-cellular ecosystems, where subclones compete for resources and cooperate to perform multiple tasks, including cancer invasion. Our recent empirical studies revealed existence of such distinct phenotypes of cancer cells, leaders and followers, in lung cancer. These two cellular subclones exchange a complex array of extracellular signals demonstrating a symbiotic relationship at the cellular level. Here, we develop a computational model of the microenvironment of the lung cancer ecosystem to explore how the interactions between subclones can advance or inhibit invasion. We found that, due to the complexity of the ecosystem, invasion may have very different dynamics characterized by the different levels of aggressiveness. By altering the signaling environment, we could alter the ecological relationship between the cell types and the overall ecosystem development. Competition between leader and follower cell populations (defined by the limited amount of resources), positive feedback within the leader cell population (controlled by the focal adhesion kinase and fibronectin signaling), and impact of the follower cells to the leaders (represented by yet undetermined proliferation signal) all had major effects on the outcome of the collective dynamics. Specifically, our analysis revealed a class of tumors (defined by the strengths of fibronectin signaling and competition) that are particularly sensitive to manipulations of the signaling environment. This class can undergo irreversible changes to the tumor ecosystem that outlast these manipulations of feedbacks and have a profound impact on invasive potential. Our study predicts a complex division of labor between cancer cell subclones and suggests new treatment strategies targeting signaling within the tumor ecosystem.

**Author Summary:** Cancer is an elusive disease due to the wide variety of cancer types and adaptability to treatment. How is this adaptability accomplished? Loss of genetic stability, a hallmark of cancer, leads to the emergence of many different types of cancer cells within a tumor. This creates a complex ecosystem where cancer cell types can cooperate, compete, and exploit each other. We have previously used an image-guided technology to isolate distinct cancer subclones and identify how they interact. Here, we have employed mathematical modeling to understand how the dynamic feedbacks between different cancer cell types can impact the success of invasion in lung cancer. We found that successful invasion required for feedbacks to support the less viable but more invasive cell types. These predictions may have implications for novel clinical treatment options and emphasize the need to visualize and probe cancer as a tumor ecosystem.

## Introduction

Lung cancer is the second most prevalent type of cancer causing over 150,000 deaths per year in the United States [1]. Insufficient progress has been made in achieving efficacious treatments. One of the main barriers in developing new treatment strategies is the vast diversity between and within cancers; heterogeneity exists between patients with the same tumor type, between tumor loci within a patient (i.e. metastases and primary tumor), and within the primary tumor itself [2,3]. Cancer is distinguished by loss of normal control over cell processes leading to genetic instability and unregulated growth. Genetic instability creates array of different clonal populations with different cell fitnesses, renewal and invasion potential [4]. Competition between different cancerous subclones and between cancerous and normal cell types sets the stage for classical ecological dynamics in the tumor microenvironment. The outcome of this process determines success of the tumor progression and its understanding may help discover novel treatment strategies [5,6].

Invasion of surrounding tissue, either locally or distally via metastasis, is a hallmark of cancer [7]. Extensive research has detailed that invasion is mediated by interactions between tumor and extracellular matrix [8,9] and cancer-associated fibroblasts [10], but there is a lack of focus on the cooperative interactions between distinct cancer subclones. Indeed, in mouse models of lung cancer, collective invasion of cancer cells was shown to correspond markedly more successful metastasis [3,11–13], confirming the critical role of collective invasion in driving cancer progression.

We recently developed a novel image-guided genomics approach termed SaGA that allowed us to identify at least two distinct phenotypic cell types in lung cancer invasion packs: highly migratory *leader cells* and highly proliferative *follower cells* [14]. Genomic and molecular interrogation of purified leader and follower cultures revealed differential gene expression prompting distinguishing phenotypes. Specifically, leader cells utilized focal adhesion kinase signaling to stimulate fibronectin remodeling and invasion. Leader cells also overexpressed many components of the vascular endothelial growth factor (VEGF) pathway facilitating recruitment of follower cells but not the leader cell motility itself [14]. However, leader cells proliferated approximately 70% slower than follower cells due to a variety of mitotic and doubling rate deficiencies. These deficiencies could be corrected by addition of cell media extracted from the follower only cell cultures, leading to conclusion that follower cells produce an unknown extracellular factor responsible for correcting mitotic deficiencies in the leader cells. In sum, leader cells provide an escape mechanism for followers, while follower cells (and follower cell media only) help leaders with increased growth. Together, these data support a service-resource mutualism during collective invasion, where at least two phenotypically distinct cell types cooperate to promote their escape.

In this new study, we developed population-level computational model to explore impact of the complex interactions between leaders and followers cell types on cancer progression. The model implemented effects of critical signaling factors controlling the communication between cell types and the interaction between cells and environment. We derived analytic boundaries dividing parameter space, representing the major signaling feedbacks, by the critical changes to invasion dynamics. Our study predicts the critical role of specific signaling pathways involved in the symbiotic interactions between cancer subclones for the overall success of cancer progression.

## Methods

Our model tracks the cell counts of leader cells, *L*, and follower cells, *F*, the concentrations of extracellular factors VEGF, *V*, an unidentified Proliferation factor, *P*, and Fibronectin, *N*, as well as the size of the domains for leader cells, *Ω*_*L*_, and for follower cells *Ω*_*F*_. Based on the available data [14], the following processes have been implemented. Leader cells can expand their domain, *Ω*_*L*_, by secreting Fibronectin, which in turn relaxes competitive pressure on leader cell growth. Leaders also secrete VEGF, which is taken up by follower cells and causes follower cells to follow them. This was modeled by increasing the domain for follower cells, which in turn relaxes competitive pressure on follower cells. Follower cells secrete an unknown proliferation signal that increases the reproductive capacity of leader cells (initially smaller than follower cells). Leader and follower cells also must compete with each other for resources at rate *c* (see Figure 1).

**Fig 1:**
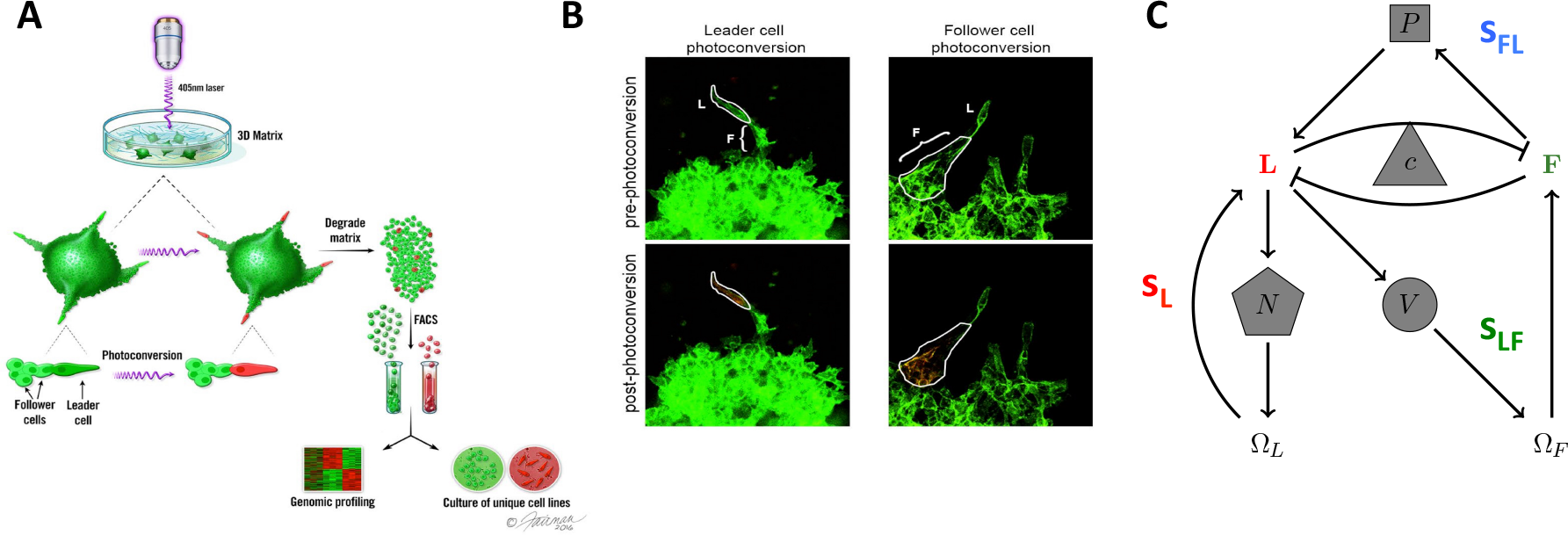
Leader and Follower system. A) Pictorial representation of the spatio-temporal genomic and cellular analysis (SaGA). Laser excitation of Dendra2 drives a change in fluorescence of user-specified cells. After degradation of cell matrix, fluorescence-based cell sorting is used to separate cells into leader and follower groups allowing for genetic analysis on specified groups. Printed with permission from Fairman Studios, LLC. This image is not included in the Creative Commons licence for the article. Adapted from Figure 1 in [14]. B) Photo-conversion examples using 3-D spheroids of H1299-Dendra2 cells. L = leader cell, F = follower cell. Adapted from Figure 1 in [14]. Stick representation of mathematical model of leader and follower cell interactions and invasion. Positive feedbacks are given by arrows, while negative feedbacks are given by flat-ended curves. The strength of leader only feedback (S_L_) is mediated by fibroNectin (N). The strength of leader to follower feedback (s_LF_) is mediated by VEGF (V). The strength of follower to leader feedback (s_FL_) is mediated by a proliferation signal secreted by followers (P). The strength of competition is given by c.

We modeled cell counts (*L* and *F* species) as standard Lotka-Volterra competition [15]. The carrying capacity of the leader cells was dynamic and dependent on the amount of proliferative signal, *P*, present. This capacity increased in a saturating manner with *P*, with maximum equal to the follower cell carrying capacity, K_*F*0_. Intra- and interspecific competition was driven by concentration, i.e. [*L*]=*L*/Ω_L_, and birthrate was driven by absolute number, *L*. The extracellular species (*V,P,N*) and domain sizes all had linear dynamics for simplicity. Below primes denote the time derivative of the variable.

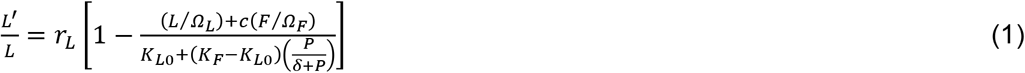

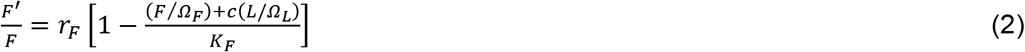

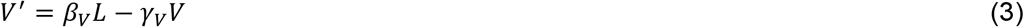

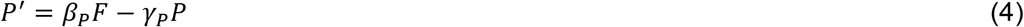

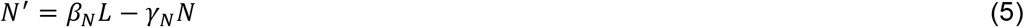

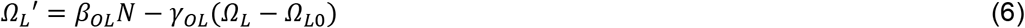

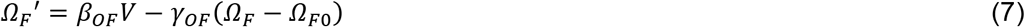

Here *r*_*L*_ and *r*_*F*_ denote the rate of expansion for leaders and followers, respectively. The parameter *c* denotes the strength of competition between the two cell types. The capacity of the environment for follower cells is given by the parameter *K*_*F*_. The capacity for leaders depended on an initial capacity, *K*_*L0*_, and on the amount of proliferation signal in a Hill-like manner with EC50, *δ*. Each extra-cellular species (*V,P,N*) had a production rate, *β*, and a degradation rate *γ*, the domain size variables (*Ω*_*L*_ and *Ω*_*F*_) also had a parameter denoting initial capacity (*Ω*_*L*0_ and *Ω*_*F*0_).

### Reduction and Feedbacks

Previous 3D spheroid experiments show that invasion occurs on a much faster time scale than reproduction [14]. By assuming that factors (*V,P,N*) and domains (*Ω*_*L*_,*Ω*_*F*_) change much faster than cell counts, one can reduce these equations to a set of two equations (*L,F*), where variables in equations (3)-(7) are at their equilibria

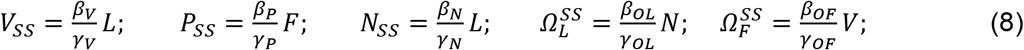

Using this reduction drastically reduced the complexity of the system. First, we defined the feedbacks based on the reduced system. The feedback that determines the leaders impact on their own domain expansion was denoted by 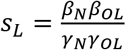, for the <u>s</u>trength of the <u>l</u>eader only feedback. The feedback that determines the leaders impact on follower cell growth was denoted by 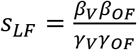, for the <u>s</u>trength of the <u>l</u>eader to <u>f</u>ollower feedback. The feedback that determines the followers impact on leader cell growth was denoted by 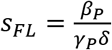, for the <u>s</u>trength of the_<u>f</u>ollower to <u>l</u>eader feedback. Second, using these assumptions, we re-wrote the leader-follower system as

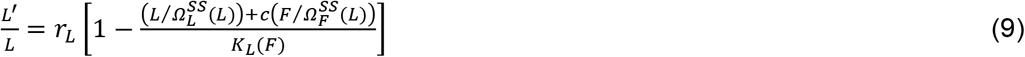

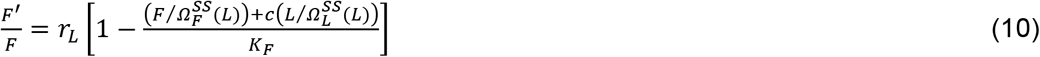

where

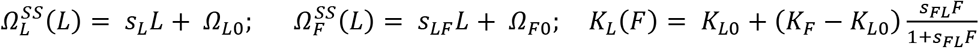

Using this reduction we can derive several critical points in invasion. The reduced system (9),(10) may have five equilibrium points: extinction of leaders (*O*_*1*_: *L*=*0*, *F*>*0*), followers (*O*_*2*_: *L*>*0*, *F*=*0*), both (*O*_*3*_: *L*=*0*, *F*=*0*), and two coexistence points (O_4_, O_5_) (where both leaders and followers populations are non-zero: *L*>*0*, *F*>*0*; *O*_*4*_ is always stable, wheras *O*_*5*_ is unstable). Changes in the feedback strengths cause fundamental shifts in dynamics. In the following we used parameter values ***Ω***_***L***_ = **1**, ***Ω***_***F***_ = **1**. To match experimental observations that leader cells grow slower and less effeciently, we set ***r***_***L***_ = **0. 3** and ***K***_***L0***_ = **0. 3** while ***r***_***F***_ = **1** and ***K***_***F***_ = **1**. The strengths of the various feedbacks, ***s***_***L***_, ***s***_***LF***_, and ***s***_***FL***_ are varied systematically below.

### Transcritical Bifurcation at Zero

To determine the critical points in the leader-follower system, we calculated the Jacobian of the reduced system evaluated for the leader extinction equilibrium (O_1_ :*L* = 0, *F* = *F*_*LE*_ = *Ω*_*F*_ · *K*_*F*_).

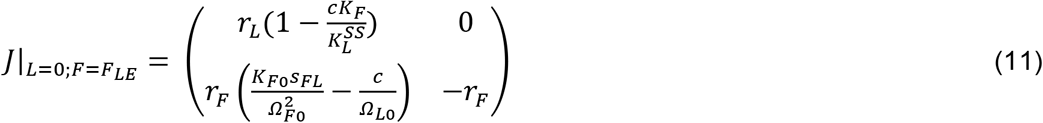

Here 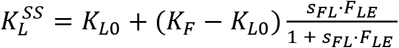, the value of *K*_*L*_ when *F* = *F*_*LE*_. The Jacobian has eigenvalues

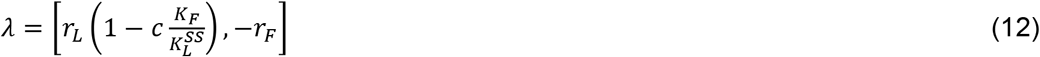

For 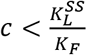, O_1_ is unstable and O_4_ (steady state where both L>0 and F>0) is stable. At 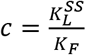 these two equilibria coincide, and for 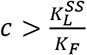 equilibrium O_4_ moves to the left of the L=0 axis and becomes unstable while O_1_ gains stability. Thus, extinction of leaders (O_1_) is stable as long as 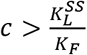, which determines an upper bound on competition where leader and followers can coexist and a bifurcation we call the transcritical bifurcation at zero.

### Saddle Node Bifurcation

The system undergoes a saddle node bifurcation when two coexistence equilibria (O_4_ and O_5_), representing non-zero populations of both leaders and followers, coincide and disapper. Beyond this bifurcation point the leader/follower populations undergo unbounded growth. This bifurcation was determined numerically using MatCont [16]. We found that this bifurcation point depends critically on both the leader feedback strength, *s*_*L*_, and on the competition strength, *c*. One of these coexistence points is effected by the transcritical bifurcation, below.

### Transcritical Bifurcation at Infinity

When the leader feedback strength is sufficiently high relative to competition, leaders and followers may undergo unbounded growth from the initial conditions belonging to the certain regions of the phase space. We describe this scenario as an attractor basin in the phase space for the stable infinity attractor. However, if *s*_*L*_ is reduced (or *c* is increased) beyond a certain threshold, infinity becomes unstable. This corresponds precisely with the loss of an unstable coexistence equilibrium with non-zero values of both leaders and followers (O_5_). Leaders and followers that are coexisting must satisfy

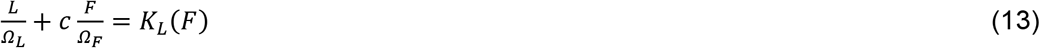

and 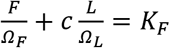 or equivalently,

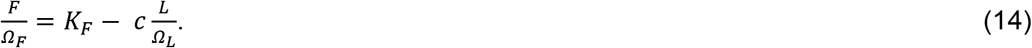

In the case that follower populations are large relative to *δ*, *K*_*L*_(*F*) → *K*_*F*_, we substituted (13) into (14) to find

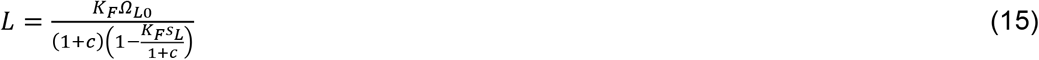

which has a discontinuity at

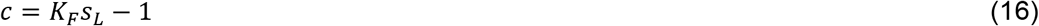

defining the loss of one of the coexistence equilibrium points (O_5_) when it moves to infinity. We describe this as the transcritical bifurcation at infinity as the stability of infinity changes at this point.

## Results

### Leader and Follower Ecosystem

Leader and follower cell types in non-small cell lung cancer spheroids were previously isolated using a fluorescence technique termed SaGA [14] (Figure 1A,B). We found that leaders and followers are genotypically and phenotypically distinct populations of cancer cells that exchange a variety of signaling molecules to coordinate complex behavior during invasion. In this new work, we focus on four main channels of communication (see Figure 1C). Leader cells secrete fibronectin in an autocrine manner. This leads to ECM restructuring and expansion of leader cell domain, Ω_*L*_, (see Methods) which ultimately increases the leader cell count. The strength of this positive feedback is characterized in our model by *s*_*L*_ (<u>s</u>trength of <u>L</u>eader only feedback). Leader cells also secrete VEGF. In the leader-follower ecosystem this promotes follower cells to track expanding leader cells, increases follower domain size (Ω_*F*_), and ultimately, follower cell count. In our model, the strength of this feedback is given by *s*_*LF*_ (<u>s</u>trength of <u>L</u>eader to <u>F</u>ollower feedback). Follower cells secrete an undetermined proliferation signal, as evidenced by the observation that follower-only cell media increases leader cell growth rate [14]. The strength of this feedback is given by *s*_*Fl*_ (<u>s</u>trength of <u>F</u>ollower to <u>L</u>eader feedback) in the model. Finally, both cell types compete for resources, which is modeled here by the feedback *c*.

These feedback mechanisms were incorporated into a modified Lotka-Volterra type competition-cooperation model. We chose a Lotka-Volterra model to focus on the ecological aspects of competition in the cancer ecosystem. Here, the leader cells could grow to a total capacity *K*_*L*_, which is an increasing function of the proliferation signal secreted by the follower cells. This capacity was reached when a combination of leader and follower cell densities (cell counts divided by domains) exceeds *K*_*L*_ (see Methods). Increases in the domain size of each type (by Fibronectin secretion in the leader case and VEGF in the follower case) limited the overall density of that cell type and mitigated its impact on the overall capacity of the system. Increasing competition, for example by limiting resources, increased the impact of either cell type on the conjugate capacity type (e.g. how leader density, L/Ω_*L*_, impacts follower capacity *K*_*F*_).

This system of the feedbacks between the leader and follower cells describes a complex dynamical ecosystem. The impact these feedbacks may have on cancer growth or invasion is unclear. Leader and follower cells are engaged in competition for resources but can also be engaged in cooperation and play supportive roles. For example, invasive leader cells provide new territory for the follower cell population and are supported by proliferative follower cells. In the following, we analyzed the model to find critical turning points for the ecosystem dynamics.

### Multiple Types of Invasion Dynamics

We found that multiple feedbacks between the leader and follower cell populations could produce a wide variety of complex dynamics. When competition strength, *c*, was high and the strength of the leader only feedback, *s*_*L*_, was moderate, population dynamic was bounded and resulted in a stable cell count for both leader and follower cell populations as well as a stable domain size (Fig. 2A). In contrast, when feedback was large and competition was moderate, population dynamics revealed an unbounded growth (Fig. 2B). Intermediate values of both *c* and *s*_*L*_ led to dynamic regimes that depended on the initial cell count: ecosystems with large initial cell count underwent unbounded growth, while small ecosystems attained a stable size (Figure 2C). These types of dynamics are in a qualitative agreement with experimental studies which revealed (a) rapid expansion of intact leader-follower ecosystem and (b) that blocking specific feedback mechanisms in vitro can reduce or block cell population growth. Specifically, blockade of fibronectin signaling or blockade of VEGF signaling led to significantly reduced invasion [14].

**Fig 2:**
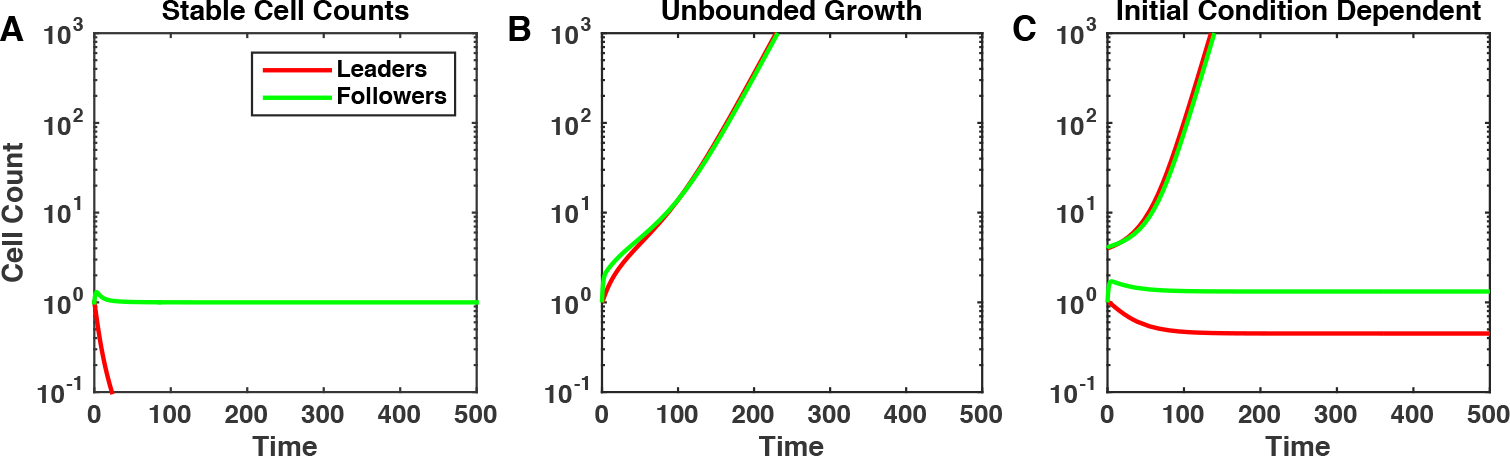
Ecosystem dynamics depend strongly on the feedbacks strength. Characteristic examples of the cancer cell population dynamics in the model for different strength of the competition between leader and followed cell populations, *c*, and leader population intrinsic feedback, *s*_L_. Cancer cell populations may attain a stable size (A: *s*_L_ = 1.2, *c* = 0.6), grow unboundedly (B: *s*_L_ = 1.2, *c* = 0.05), or be dependent on the initial tumor size (C: *s*_L_ = 2, *c* = 0.375).

This array of behaviors can be explained by the critical shifts in the cell population dynamics due to the changes in the feedbacks strength. We found that depending on the level of competition, *c*, and the strength of invasiveness of leaders, *s*_*L*_, the leader-follower ecosystem can operate in one of five different regimes, as described below (Figure 3).

**Fig 3:**
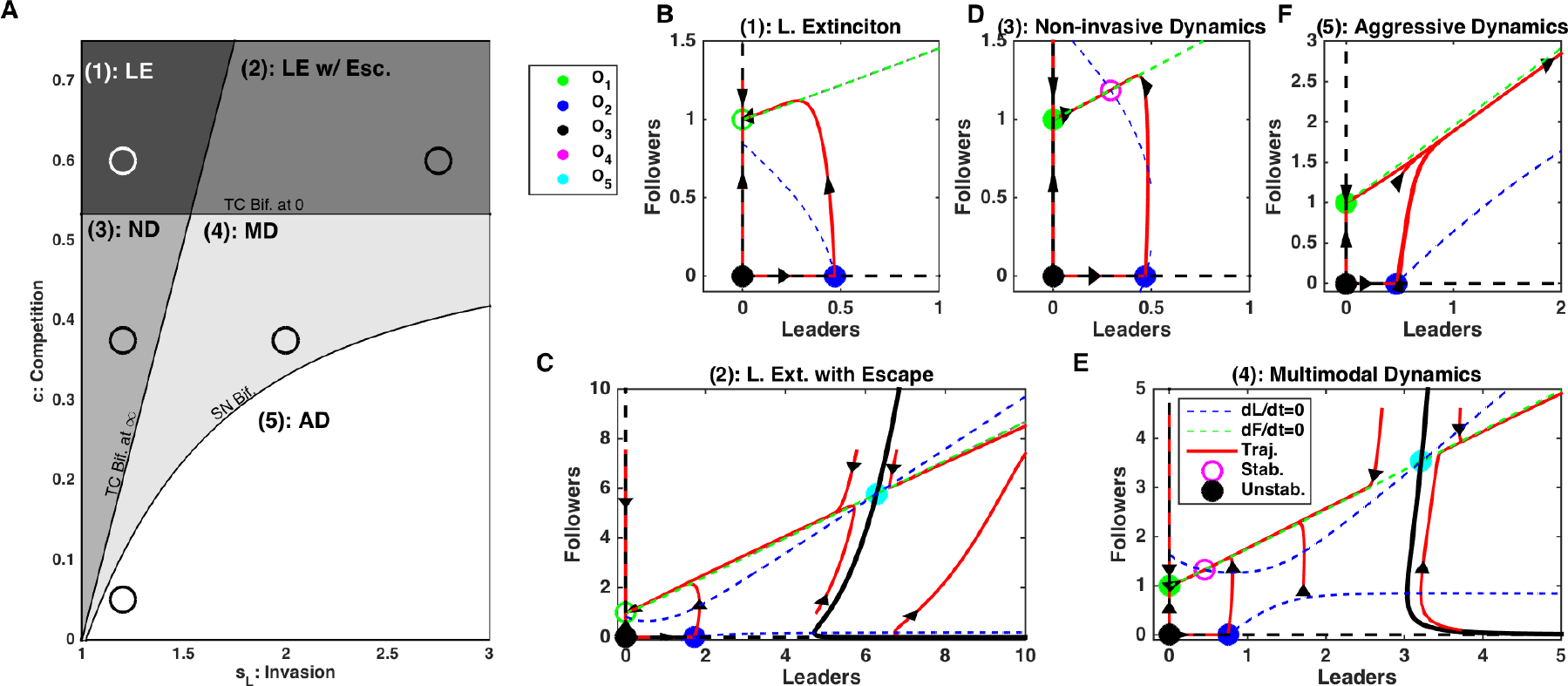
Dynamics of the leader and follower ecosystem. A) Bifurcation diagram sweeping leader population intrinsic feedback strengths, *S*_*L*_, and competition strength, *c*. Abbreviations - LE: Leader Extinction, LE w/ Esc.: Leader Extinction with Escape, ND: Non-invasive Dynamics, MD: Multimodal Dynamics, AD: Aggressive Dynamics, TC Bif. at ∞: Transcritical Bifurcation at infinity (see Methods), TC Bif. at 0: Transcritical Bifurcation at zero (see Methods), SN Bif.: Saddle Node Bifurcation (see Methods). B-E) Phase diagrams for each regime identified in A). Dashed blue and green lines are null-clines for the leaders and followers, respectively. Red curves show the trajectories of the leader-follower system from different initial conditions, with arrows denoting tangent vectors at various points. Open circles denote stable equilibrium points, stars denote unstable equilibria. Specific *s*_*L*_ and *c* parameters used in panels B-E) are given by the circles in A).

**Leader Extinction:** When competition was high and invasive feedback was minimal, the leader cells (the weaker competitor) were forced to extinction while the follower cells persisted and its population reached a stable size (Figure 3A,B). There was a critical level of competition between leaders and followers, given by 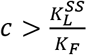 (see Methods, *Transcritical bifurcation at zero* for derivation), required for this type of dynamics. This critical level of competition, the ratio of the capacity of leader cells, 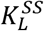, to that of the follower cells, *K*_*F*_, essentially depends on the fitness differences between leader only and follower only cell populations. Leader and follower populations with similar fitness would tolerate a much higher competition threshold without driving one species to extinction. From the dynamical systems perspective, when 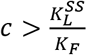 and *s*_*L*_ is sufficiently low (see below), the only stable equilibrium in the phase space is O_1_ and all system trajectories converge to this equilibrium point representing the leader cells extinction state (Figure 3B).

**Leader Extinction with Escape:** If competition was above the leader extinction limit, 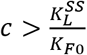, but not high enough to balance the impact of the leader only feedback, *c* < *K*_*F*_*s*_*L*_ − 1, there were two possible outcomes depending on the initial population size (see Methods, *Transcritical Bifurcation at Infinity* for derivation) (Figure 3A,C). The second condition, *c* < *K*_*F*_*s*_*L*_ − 1, can be interpreted as a balance between positive feedback, *s*_*L*_, and negative feedback, *c*. In this regime, leaders could go extinct if the initial population of leader cells was sufficiently small. Alternatively, if initial populations of leaders and followers both were large enough, the ecosystem could grow unboundedly. Thus, our model predicts, that the ability to undergo successful collective invasion depends on whether the initial bulk size is larger than a critical amount. These types of dynamics with divergent outcomes occur when competition is large enough to be able to drive leaders extinct, but small enough so it can be outbalanced by the strong invasive effects of the leader cells.

In the phase space of the model, the basins of attraction of the two distinct dynamical regimes are separated by a critical boundary (separatrix of a saddle equilibrium O_5_) where the cell bulk size determines its ultimate fate (Figure 3C). Both infinity and the leader extinction equilibrium (O_1_) are stable attractors representing two possible end solutions of the system dynamics.

**Non-invasive Dynamics:** When competition was smaller than the extinction limit, 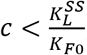 but large enough to balance leader feedback strength, *c* > *K*_*F*_*s*_*L*_ − 1, the ecosystem size remained bounded and both leaders and followers attained a stable and non-zero population size. In the phase space, this type of dynamics corresponds to conversion to the stable equilibrium O_4_ (Figure 3A,D). We refer to this as non-invasive dynamics, as the cells cannot grow beyond a defined size. In this case, while competition was present, it was too weak to lead to extinction, while leader population was not invasive enough to promote unlimited growth. This scenario represents stable, non-invasive dynamics.

**Multimodal Dynamics:** If competition was (a) small enough to allow leader existence, 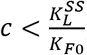 (b) small enough relative to the leader feedback strength, so that escape was possible, *c* < *K*_*F*_*s*_*L*_ − 1, but (c) high enough, so that for small initial population of leader it could balance the positive leader feedback, leader and follower cell dynamics depended on the initial population size (Figure 3A,E). Ecosystems with a large initial cell count would grow without bound but those with a small initial cell count would reach a stable population size, due to the competition as in the non-invasive dynamics case. On the phase plane the last outcome was represented by a contraction to a stable equilibrium O_4_. This critical boundary was defined by a separatrix of a saddle fixed point O_5_ (Fig. 3E). (This separatrix was determined numerically by reversing time [17].)

**Aggressive Dynamics:** When leader invasive strength was sufficiently high and competition was sufficiently low, the only possible outcome was unbounded growth of both cell populations (Figure 3A, F). In this case, the only stable attractor in the phase space is infinity where all system trajectories are converged to.

In summary, our analysis revealed that the complex balance of the feedbacks in the leader-follower ecosystem can lead to the multiple types of population dynamics. When the leaders’ invasiveness was low, the outcome depended on the competition between two populations – strong enough competition promoted leader extinction, while weak competition allowed stable coexistence states with bounded size of both leader and follower cell populations. As leader invasiveness rate increased, the system revealed a new state with unbounded growth. This aggressive dynamic state coexisted with a stable attractor representing a bounded size of both populations if competition between leaders and followers was strong enough. Otherwise, unlimited population growth was the only outcome. Based on the system dynamics derived above, next we will show how critical boundaries between parameter regimes could be exploited to lead to profound changes in the ecosystem dynamics.

### Limiting leader feedback Leads to Irreversible Changes in Invasion

In the multimodal dynamics (e.g., Fig. 3C or 3E), leader and follower cell populations can undergo explosive growth or achieve a stable count depending on the initial size of the ecosystem. We examined the impact of limiting the invasive leader feedback in scenarios of this type (Fig. 4). Even when the ecosystem was initially sufficiently large to support unbounded growth, after reducing invasive leader feedback *s*_*L*_ (Fig. 4A), the ecosystem was forced into the non-invasive dynamics type and the total bulk of the cell population reduced reaching a steady-state (Fig. 4E). Importantly, the leader and follower cell populations remained stable and bounded after restoring invasive leader feedback to its original strength (Fig 4E, right side).

**Fig 4:**
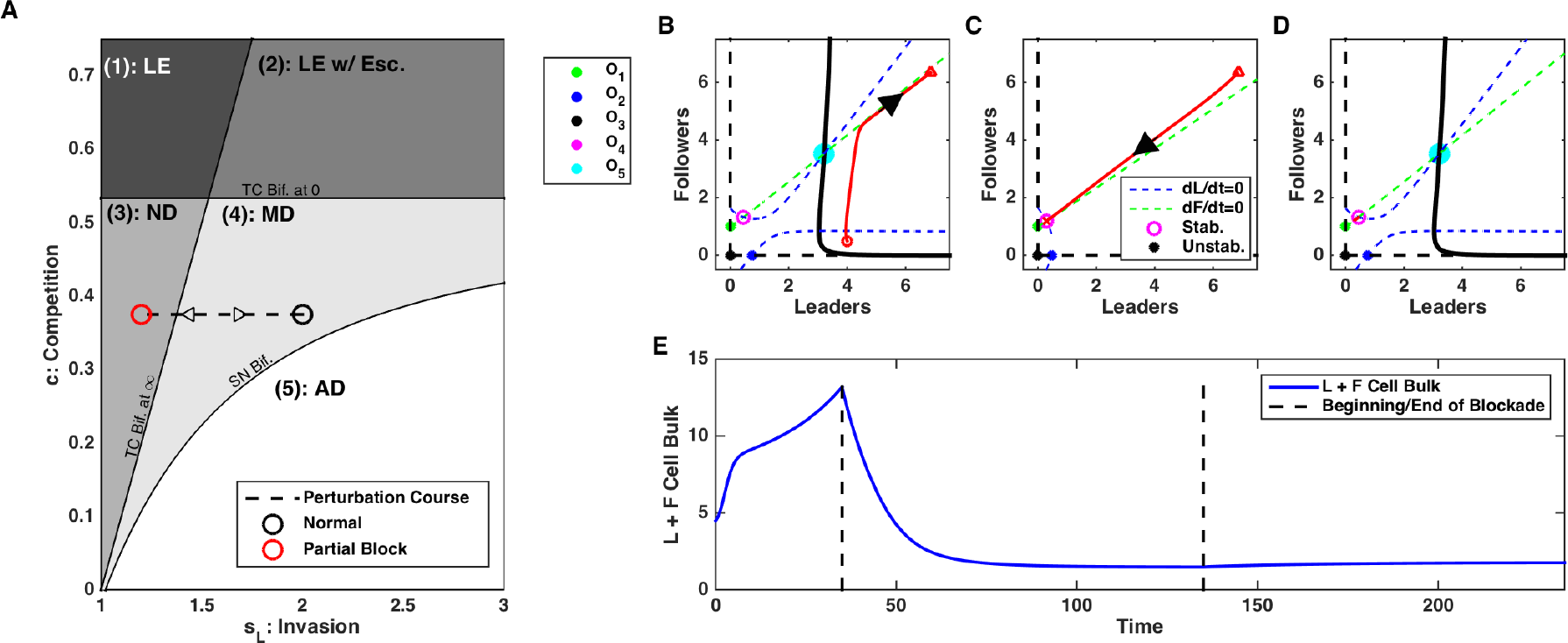
Blocking leader cell population feedback in the Multi-modal Dynamics regime can lead to irreversible changes of the population dynamics. A) Bifurcation diagram depicting the direction of the perturbation in the parameter space - increase in feedback, s_L_. Black circuit - initial state; red circuit - perturbed state. B-D) Phase plots of dynamics before (B), during (C), and after (D) blockade of *s*_*L*_. E) Time-course of cell bulk before, during, and after perturbation. Note irreversible change of the ecosystem dynamics.

From the point of view of the dynamical systems analysis, reducing leader feedback changed the phase space, so the only stable attractor was non-zero equilibrium (O_4_) (Fig. 4C). In this regime, unlimited growth was abandoned and the system converged to the equilibrium state (O_4_) corresponding to the bounded size of both cell populations. This equilibrium remained stable even after the feedback was restored to its original level (Fig. 4D).

Our model predictions (Fig. 5A) are consistent with in vitro data (Fig. 5B). Using siRNA blocking we previously showed that expression of fibronectin (which is characterized by the <u>s</u>trength of <u>l</u>eader only feedback, *s*_*L*_, in the model) led to the low invasion potential and a stable cell population size [14].

**Fig 5:**
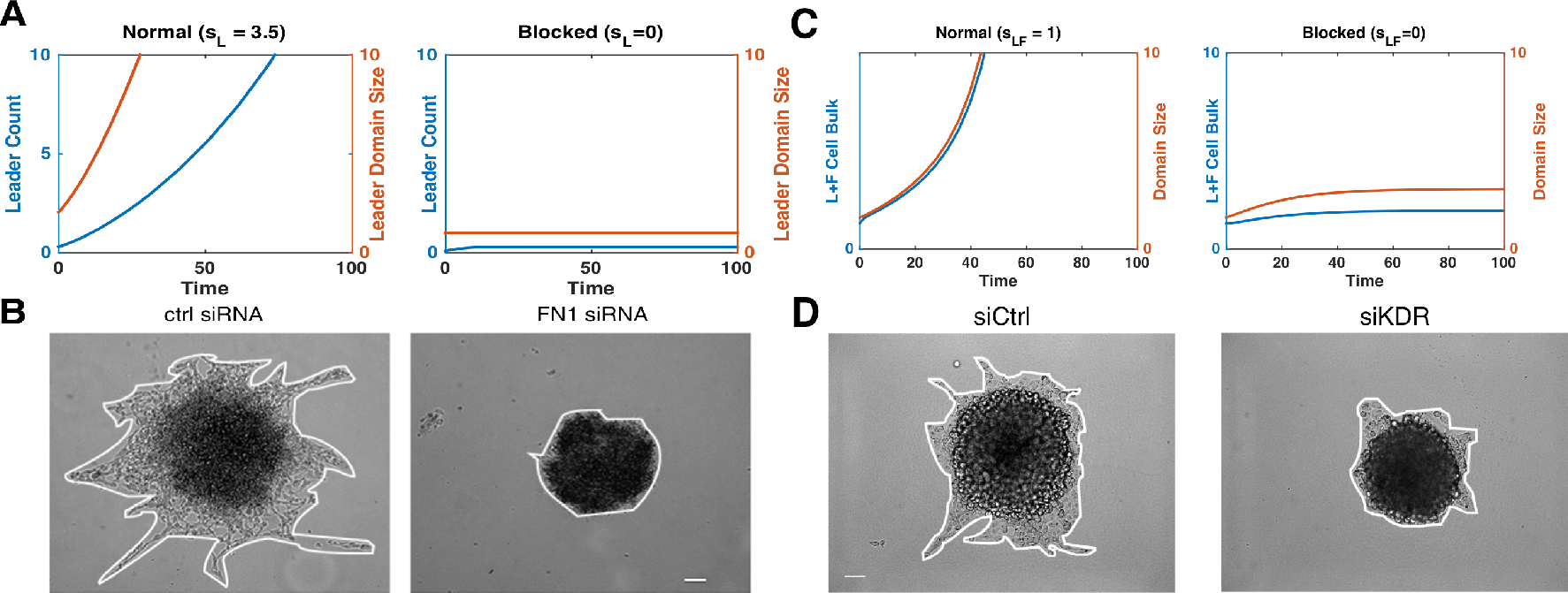
Model reproduces in vitro experimental results. A) In the model, leader cell count (blue) and domain size (red) are given in both normal (left) and leader population feedback, s_L_, blocked (right) conditions. B) In cell culture, invasion of leader cells was significantly reduced during siRNA block of focal adhesion kinase (right), compare to control (left). Scale bar 100*μ*m. Reproduced from [14]. C) In the model, blocking leader to follower feedback, s_LF_, limited invasive area and cell count. D) Impact on invasion of leader cell cultures during siRNA block of VEGFR2 (siKDR). Scale bar 100*μ*m. Invasive area was significantly reduced after VEGFR2 block (p<0.0001) (right), compare to control (left). Reproduced from [14].

### Increasing Competitive Signals Leads to Leader Extinction

We next tested effect of increasing competition between leader and follower cell populations on the ecosystem dynamics (Fig. 6). Leader cells excrete extracellular factors that induce the death of the followers and leaders alike [14], which supports competition. Here, we again started from aggressive unbounded type of dynamics and then increased competition strength (Fig. 6A). This caused change of the ecosystem dynamics. Both cell populations reduced the size, with leader cell population going to extinction state (Fig. 6E). However, upon restoring competition to the original level, leader and follower cells reemerge and grow unboundedly again. The last can be avoided if no leader cells remain (complete extinction).

**Fig 6:**
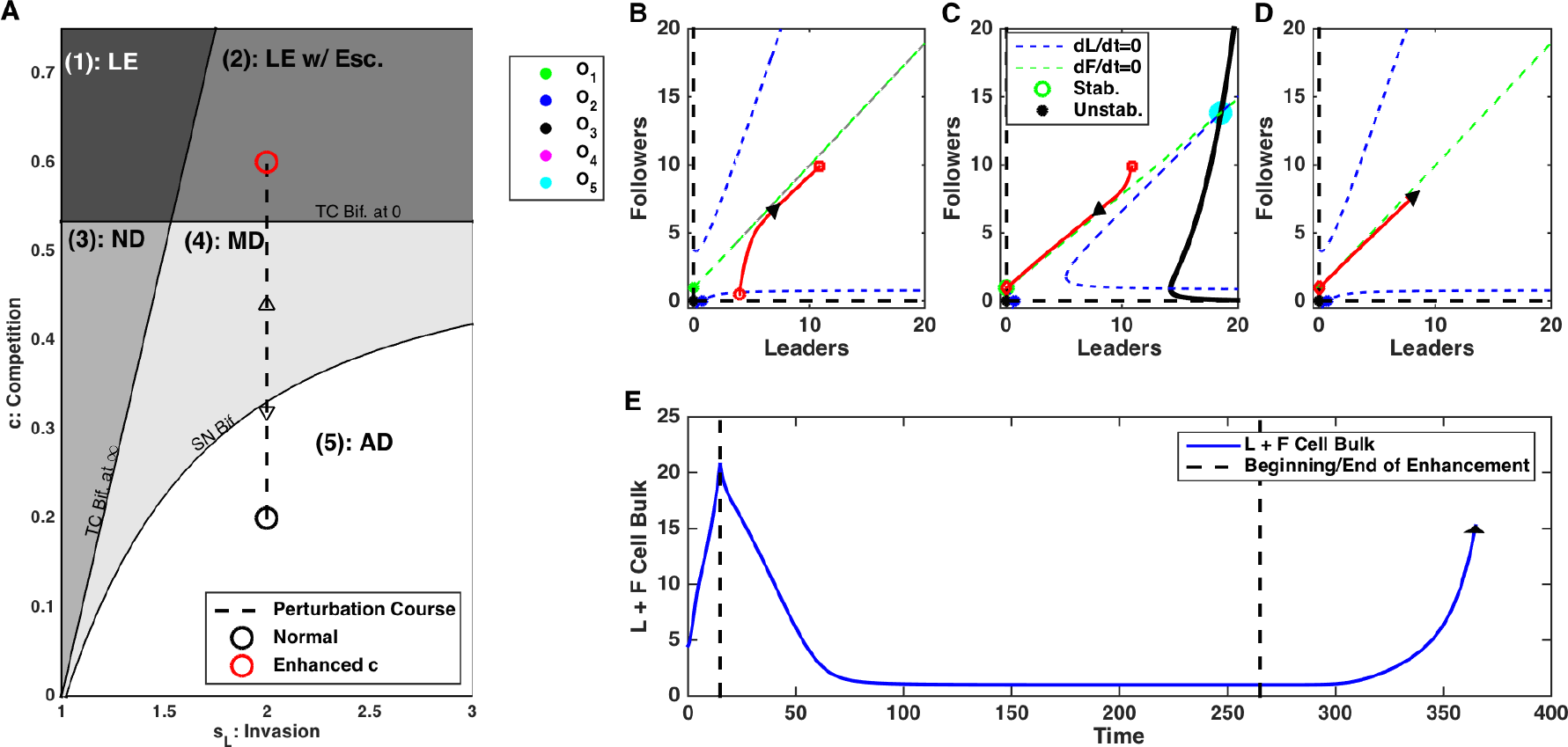
Enhancing competition between leader and follower populations can drive transient extinction of leaders. A) Bifurcation diagram depicting the direction of the perturbation - increase in competition, *c*. Black circuit - initial state; red circuit - perturbed state. B-D) Phase plots of dynamics before (B), during (C), and after (D) enhancement of competition. E) Time-course of cell count. Here, we assume total extinction of leaders occurs during treatment, i.e. at some point during treatment *L*=*0*. Note that ecosystem dynamics is reversed after perturbation is removed.

Again, this dynamic can be easily understood using bifurcation analysis. Increasing competition strength made leader extinction equilibrium state O_1_ stable (Fig. 6C). However, when competition was restored to its original level, O_1_ became unstable again and leader and follower cells returned to escape dynamics (Fig. 6D). Importantly, in the extreme case of very small cell populations, cells undergo discrete and stochastic dynamics and complete extinction of a small population of leaders is possible in a finite time, leading to irreversible changes due to competition increase (similar dynamics was described in our previous study [18]).

### Support For Leaders has Large Impact on Aggressiveness

Changing the strength of the feedbacks that determine the interaction between leaders and followers (s_LF_ and s_FL_) could also impact the dynamics. Leader cells secrete VEGF (denoted here by s_LF_) that helps follower cells to expand their territory and follower cells secrete a proliferation signal (denoted here by s_FL_) that allows leaders to increase their proliferative capacity. These two feedbacks have distinct impacts on the overall ecosystem dynamics. Perturbations to s_LF_ (changing the impact that leaders have on followers) changed the system dynamics (assuming that the cell count was small enough at the time of the intervention) from unlimited growth to the bounded type. The size of both leader and follower cell populations decreased reaching non-zero steady-state (Fig. 7E). This regime persisted as long as the feedback from the leaders to followers remained low. However, increasing s_LF_ to its original level restored the system dynamics with unlimited cell population growth (Fig. 7E, right size).

**Fig 7:**
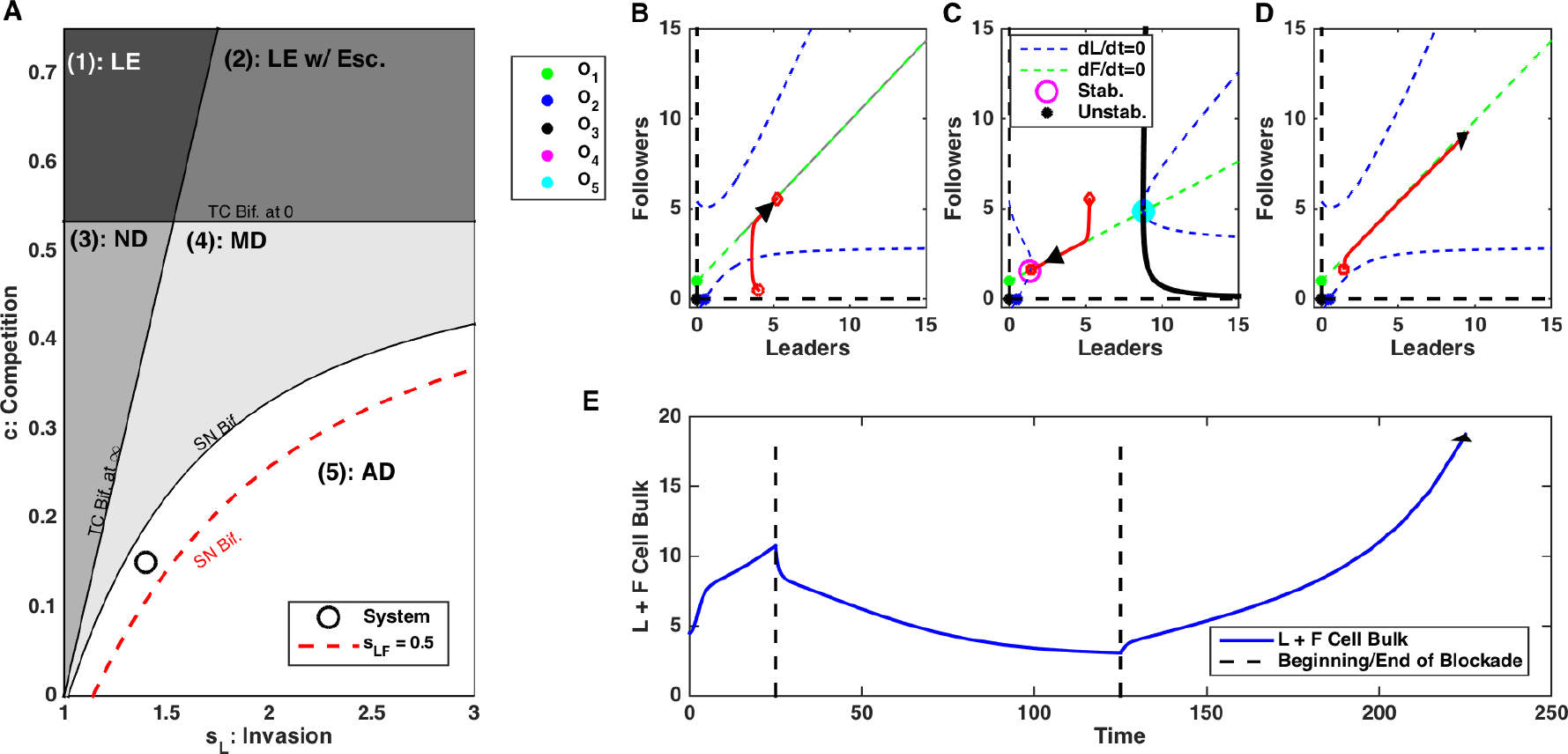
Disrupting Leader to Follower feedback, s_LF_, can trigger transient changes in the population dynamics. A) Bifurcation diagram depicting the direction of the perturbation in parameter space. Perturbations in *s*_*LF*_ change the location of the saddle node bifurcation boundary. Black line - initial location of the bifurcation boundary; red line - perturbed location. B-D) Phase plots of dynamics before (B), during (C), and after (D) blockade of *s*_*LF*_. E) Time-course of cell count. Note that ecosystem dynamics is reversed after perturbation is removed

Using bifurcation analysis, we found that reducing impact that leaders have on followers shifted the location of the saddle node bifurcation boundary that separated state with unlimited growth only dynamics and a state with coexistence of the unlimited growth and a stable equilibrium attractor (O_4_) regimes (Fig. 7A). Effectively, decreasing sLF increased the threshold level of the invasive leader feedback (s_L_) needed to cause unbounded growth. Thus, reducing s_LF_ made the system to converge to the stable equilibrium state O_4_ corresponding to the bounded size of both cell populations (Fig. 7C). However, increasing s_LF_ to its original level changed the phase space again, so infinity became the only stable attractor (Fig. 7D) and unlimited growth dynamics resumed.

Our model predictions (Fig. 5C) are consistent with in vitro data (Fig. 5D). Using siRNA to block the VEGF receptor VEGFR2 (siKDR in Fig. 5D), we previously showed that blocking the leader to follower feedback led to the limited invasion potential and stable cell population size (Fig. 5D) [14].

Finally, we tested the role of the follower to leader feedback (s_FL_) and found that perturbations to s_FL_ have a significant impact on the system dynamics. In contrast to s_LF_, changes to the s_FL_ changed both the location of the saddle node bifurcation boundary and the transcritical bifurcation boundary of the leader extinction (Fig. 8A). Therefore, decreasing s_FL_ both increased the threshold on the leader invasion strength (s_L_) needed to cause unbounded population growth and decreased the threshold of the competition strength (c) needed to induce leader population extinction. We have exploited this to show that decreasing s_FL_ can cause irreversible change in the cell population bulk. Again, starting with unlimited growth dynamics (Fig. 8B), decreasing follower to leader feedback, s_LF_, reversed the dynamics and both leader and follower cell population reduced in size converging to the steady-state (Fig. 8E). This regime with bounded ecosystem size persisted after the feedback was restored (Fig. 8E, right side).

**Fig 8:**
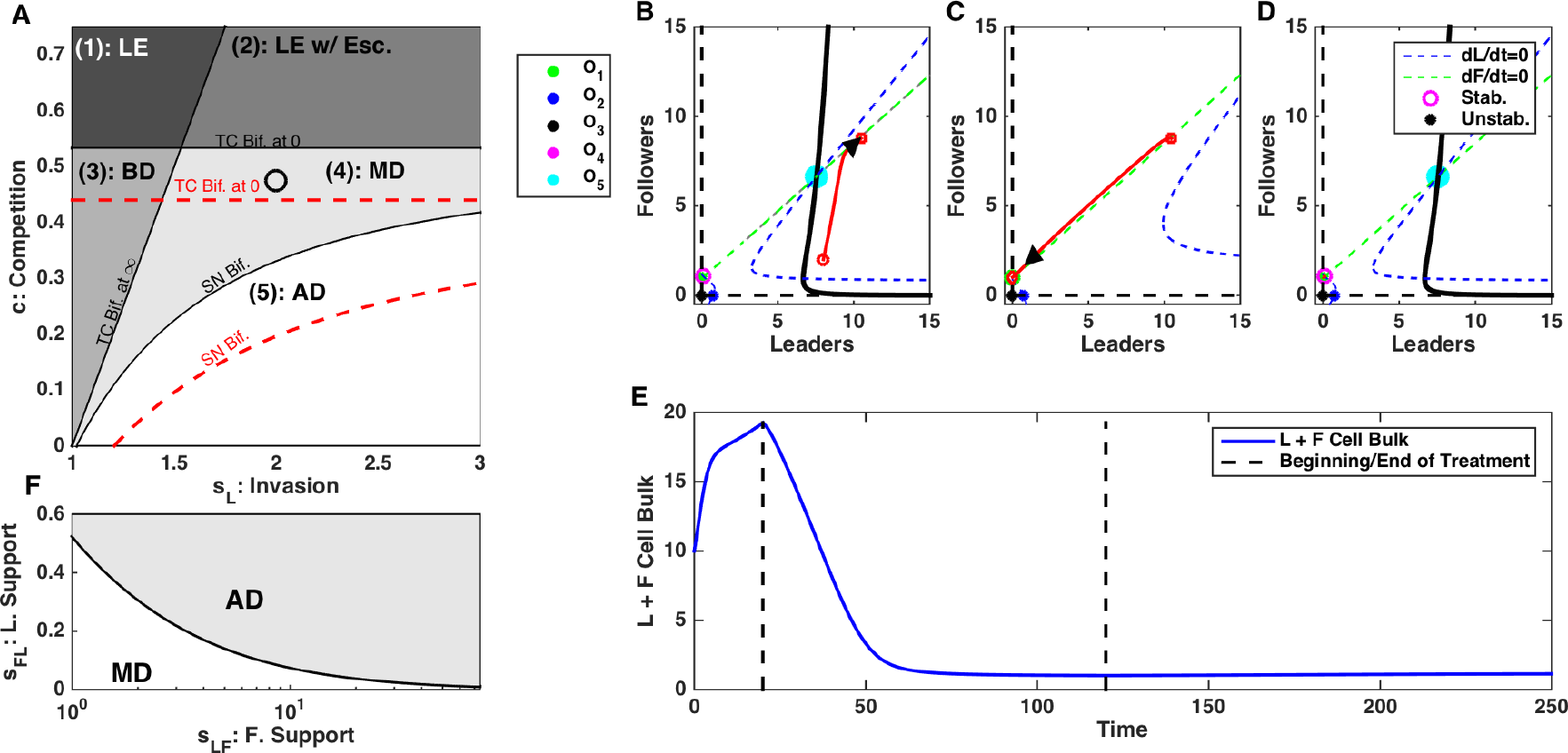
Disrupting Follower to Leader feedback, s_FL_, can have irreversible changes in dynamics leading to stabilization of cell count. A) Bifurcation diagram depicting the direction of the perturbation in parameter space. Perturbations in *s*_*FL*_ change the location of the saddle node bifurcation and transcritical bifurcation at zero boundaries. Black lines - initial location of the bifurcation boundaries; red lines - perturbed location. B-D) Phase plots of dynamics before (B), during (C), and after (D) blockade of *s*_*FL*_. E) Time-course of cell count. Note irreversible change of the ecosystem dynamics. F) Bifurcation diagram depicting the position of the saddle node bifurcation point as a function of *s*_*FL*_ and *s*_*FL*_. AD: Aggressive Dynamics; MD: Multi-modal Dynamics. Here, *s*_*L*_ = 1.4 and *c* = 0.2.

Using dynamical systems analysis, we found that reducing follower to leader feedback (s_FL_) triggered the system convergence to the stable attractor (O_1_) representing the leader extinction state (Fig. 8C). When the feedback was restored, O_1_ becomes unstable but the ecosystem fell to the attraction basin of the stable equilibrium O_4_ and avoided regime of unlimited growth (Fig. 8D). In more general case, the outcome depended on the balance between the leader to follower, s_FL_, and follower to leader, s_LF_, feedbacks, with higher s_LF_ requiring more significant s_FL_ decrease to avoid unbounded growth (Fig. 8F).

### Summary of Perturbations to Cancer Ecosystem

Complex balance of the feedbacks within the cancer cell ecosystem allows for some alterations of the feedback parameters to have significant impacts on the ecosystem dynamics. We summarized these different possibilities in Table 1 from the perspective of achieving the goal to reduce cell population bulk. Hence, manipulating *s*_*L*_, *s*_*LF*_, *s*_*FL*_ should be interpreted as decreasing these feedbacks, whereas manipulating *c* should be interpreted as increasing *c*. We also examined the possibility of non-targeted cell death, such as might occur during non-specific chemotherapy. Manipulations were either irreversible, so the system dynamics remained altered upon cessation of the perturbation (e.g. irreversible leader extinction or irreversible stabilization of the cell count), or caused only temporal and reversible reduction of the cell bulk. In some cases, such as leader extinction with escape and multimodal dynamics (see Fig. 3), the size of the initial cell bulk dictated possible outcomes of the feedback perturbations. The outcomes described in the Table 1 represent the best-case scenario. Thus, perturbations were started from an appropriate initial state and maintained long enough to achieve the desired effect.

**Table 1:** Effect of the different feedback alterations on the ecosystem dynamics. First column - the different types of the ecosystem dynamics in the (s_L_, c) parameter space are as shown in Figure 3. In some cases, initial population size dictated the outcome. These outcomes are distinguished in the initial state given in the 2d column. An initial state in the infinity attractor basin denotes that the cell bulk exceeded the critical amount and could grow unboundedly; initial state in the stable attractor basin denotes the case when the cell counts were less than the critical value and the system converged to the stable attractor. Third column - each transient manipulation to the model parameters had a goal to reduce cell population size (decrease of *s*_*L*_, *s*_*LF*_, *s*_*FL*_ or increase of *c*). The “death” indicates a non-targeted “enforced” reduction of the cell population. The last column indicates the system dynamics after the original values of the model parameters were restored. The changes were either irreversible upon cessation of the perturbation (e.g. irreversible leader extinction and irreversible stabilization of cell bulk) or caused reversible reduction in the cell population bulk.

| Type of dynamics | Initial State | Manipulation Type | Outcome |
| --- | --- | --- | --- |
| 1. Leader Extinction | N/A | death, $c$, $s_L$, $s_{LF}$, $s_{FL}$ | Reversible cell bulk reduction. |
| 2. Leader Extinction w/ Escape | Infinity attractor basin<br>Stable attractor basin | death, $s_L$, $c$, $s_{LF}$, $s_{FL}$<br>death, $c$, $s_L$, $s_{LF}$, $s_{FL}$ | Irreversible leader extinction.<br>Reversible cell bulk reduction. |
| 3. Non-invasive Dynamics | N/A | $c$, $s_{FL}$, $s_L$, $s_{LF}$, death | Reversible cell bulk reduction. |
| 4. Multimodal | Infinity attractor basin<br>Stable attractor basin | $c$, $s_{FL}$, $s_L$, $s_{LF}$, death<br>$c$, $s_{FL}$, $s_L$, $s_{LF}$, death | Irreversible stabilization of cell bulk.<br>Reversible cell bulk reduction. |
| 5. Aggressive Dynamics | N/A | $c$, $s_{FL}$, $s_L$, $s_{LF}$, death | Reversible cell bulk reduction. |

This analysis revealed that certain parameter regimes are more sensitive to the perturbations than others. Specifically, in the leader extinction with escape regime (area (2) in Figure 3A) and the multimodal dynamics regime (area (4) in Figure 3A) perturbations could have irreversible impacts on the ecosystem. In these cases, any perturbation (death, reduction in *s*_*L*_, *s*_*LF*_, *s*_*FL*_, or increase in *c*) can potentially force the system to cross the critical boundary (separatrix) and transition from explosive growth to a steady-state dynamic. These regimes give a unique opportunity to impact the invasiveness of the ecosystem.

Also, certain perturbations could force the ecosystem into a state where leader extinction (*O*_*1*_) is stable. This occurs when applying sufficient increases in the competition pressure, *c*, or decreases in the support from followers to leaders, *s*_*LF*_. In these cases, it is possible for the discrete and stochastic nature of the cell population dynamics to define the ecosystem fate. Thus, a sufficiently long perturbation could irreversibly eradicate a sufficiently small discrete number of leader cells [18].

## Discussion

Heterogeneity of tumors, at the genetic, epigenetic, and phenotypic levels, is one of the main obstacles to developing new effective treatment strategies. Tumor cells rapidly evolve forming highly efficient symbiotic systems with well-defined labor division targeted to augment tumor survival and expansion. In lung cancer collective invasion packs observed *in vitro*, two distinct populations of cancer cells - highly migratory *leader cells* and highly proliferative *follower cells* - have been recently identified [14]. In this new study, we used computational models to explore collective dynamics of the leader-follower ecosystem and to exploit approaches that can effectively disrupt it.

We found that competition between two populations (defined by the limited amount of resources), the positive feedback within the leader cell population (controlled by the focal adhesion kinase and fibronectin signaling) and impact of the follower cells to the leaders (represented by yet undetermined proliferation signal) all had major effects on the outcome of the collective dynamics. While increase of the positive feedback within the leader cell population would ultimately lead to the system state with unbounded growth, manipulating follower to leader feedback or increasing competition between leader and follower cell populations was able to reverse this dynamic and to form a stable configuration of the leader and follower cell populations.

Our model highlights the importance of focal adhesion kinase (FAK) and fibronectin signaling. Our previous empirical work showed that FAK signaling was a key distinguishing feature between leader and follower cells and critical for invasive leader behavior [14]. Our model predicts that FAK is the main driver of invasion by leader cells and disruptions in the FAK driven feedback loop cause critical changes in the leader-follower population dynamics. Indeed, FAK is a well-known regulator of the tumor microenvironment: promoting cell motility and invasion [19]. FAK expression is upregulated in ovarian [20] and breast cancer [21] tumors with expression levels correlating with survival [22,23]. Many FAK inhibitors, such as defactinib, are currently in clinical trials with promising results [19],24,24–28]. A key advantage of FAK inhibitors is that they impact both the tumor itself and the surrounding stroma where tumor associated fibroblasts also utilize FAK signaling to promote tumor invasiveness [29,30].

While commonly associated with angiogenesis in healthy and cancerous tissue, our previous work showed that VEGF mediates communication between leader and follower cells [14]. There is a long history of targeting VEGF to limit tumor invasiveness [31,32]. While great success has been seen in preclinical models [33,34], only moderate success was seen in clinical trials with anti-VEGF drugs such as bevacizumab [35,36]. This is largely due to cancers developing resistance to specific VEGF-therapeutics. In our model, VEGF stimulated followers to shadow leaders and expand their domain. However, we found that inhibition of VEGF had little impact on the ecosystem dynamics relative to the perturbations of the other axes (such as FAK or competition for resources).

Competition for resources is one of the principal forces that structures any ecosystem, including tumor ecosystems [6,37]. Our modeling work predicts that competition was a critical component in the leader-follower ecosystem. We found that when the strength of competition exceeded a critical threshold, leaders (the weaker competitor) were driven to extinction. Further, enhancements of the competition in the model changed the fundamental cell population dynamics. In some cases this meant stopping unbounded growth and promoting the extinction of the leader cells. Our previous in vitro work demonstrated that leaders may inhibit the growth of followers through an unknown secreted factor in cell media [14]. While still in the early stages, exploiting this inhibition may also provide similar benefits to those shown here as increases in competition.

Our previous study also revealed a currently unknown extracellular factor secreted by followers that corrects mitotic deficiencies and enhances leader proliferation [14]. Our modeling highlights this factor as having critical impact on the ecosystem dynamics. We found that blockade of this proliferation factor, modeled here by the strength of the followed to leader feedback, can cause critical shifts in the population dynamics. More work needs to be done to identify and understand the mechanism of this action, but preliminary results suggests that this may be a potential novel treatment axis that specifically targets the mutualistic interaction between leaders and followers.

Ecological forces shape the exchange of biomaterial between different biotic and abiotic environmental agents. These forces determine capacity of the ecosystem for different species (subclones) and the environment ultimately sets the fitness of each of the competitors. Classic ecological theory dictates that an abundance of many similar species (such as similar subclonal populations) will lead to a high competition for resources [38,39]. This competition can force the exclusion of inferior competitors and ultimately may reduce heterogeneity of the system. However, when symbiotic and mutualistic interactions occur, otherwise competitive species support each other and increase the capacity of the ecosystem [40,41]. Symbiosis between different subclonal populations may be particularly important during critical times when the tumor survival is in peril (such as hypoxia, metastasis or therapy). One critical moment in tumor progression occurs when highly proliferative tumor cells saturate the resource potential of their current environment. In order to obtain more resources, tumors need to invade new territory.

Previous results to model complex tumor cell population dynamics range from detailed cellular level models (e.g. [9,42–44]) to continuous models with a different degree of complexity (e.g. [18,45–48]) similar to that proposed in our new study. While cellular level model can directly incorporate heterogeneous cell types and intrinsic tumor properties, including proliferation, metabolism, migration, protease and basement membrane protein expression, and cell-cell adhesion, they typically have highdimensional variables and parameter space that is difficult to explore. Advantages of the reduced type of models include the low dimensional parameter space, where parameters have clear biophysical meanings, and which allows for systematic analysis to rapidly explore and determine the sensitive parameter space. We previously applied this approach to study cell interactions in chronic cancers and predicted conditions for explosive tumor growth [18]. Similar approach was applied to model cancer cell population dynamics in many other types of cancer [45,48,49].

The vast diversity between different cancers and even between different cell types within a single tumor remains one of the biggest hurdles to overcome to achieve personalized cancer treatment. This diversity leads to a complex array of interactions between different tumor cell types and the healthy surrounding tissue: the tumor ecosystem. Our work has isolated phenotypically unique lung cancer cells and taken a dynamical approach to understanding the interactions within the tumor ecosystem. We identified the critical features and interactions composing the leader-follower ecosystem, to explore vulnerabilities of the lung cancer invasive cell populations.

